# Comparative Distribution of α2,3- and α2,6-Linked Sialic Acid Receptors and Influenza A(H5N1) Virus Binding in Sheep, Goats, and Alpacas

**DOI:** 10.64898/2026.10.26.762033

**Authors:** Nayrmaliz Marrero, Eli Blay, Serban Oancea, Gulnara Brixius, Rahul Nelli, Paul J. Plummer, Jennifer Groeltz-Thrush, Breigha Boyle, Brianna Jones, Olufemi Fasina, Tyler Harm, Chris Siepker, Rachel Phillips, Laura Bradner, Alissa Hufnagel, Michael Zeller, Eric Cassmann, Silvia Carnaccini, Todd M. Bell, C. Joaquin Caceres

## Abstract

Highly pathogenic avian influenza virus (HPAIV) H5N1 continues to pose a significant threat to animal and human health. Since 2022, an expanded host range has been observed, as illustrated by the H5N1 outbreak in dairy cows. Other minor livestock species, such as sheep, goats, and alpacas, have also tested positive for H5N1 for the first time, raising concerns about its impact on previously overlooked livestock species. In this study, we first characterized the distribution of sialic acid (SA) receptors across respiratory and non-respiratory tissues of sheep, goats, and alpacas using *Maackia amurensis* (SA α2,3) and *Sambucus nigra* (SA α2,6) lectins. We found that both SA receptors are present in all tissues across species, with notable differences in their distribution and abundance. Next, we developed recombinant LPAIV H5N1 viruses over a laboratory-adapted backbone to evaluate tissue-specific virus binding profiles. The results show that a representative H5N1 virus from clade 2.3.4.4b binds effectively to respiratory and non-respiratory tissues across all species. The strongest virus binding was observed in sheep, followed by goats and, lastly, alpacas. Similar binding patterns were observed with an LPAIV version of clade 2.3.4.4b and a Gs/Gd lineage precursor prototype virus. Finally, we show that H1N1 and H3N2 viruses also bind to respiratory tissues of sheep and goats. These findings provide important insights into the potential susceptibility of minor livestock species to HPAIV H5N1 and other influenza subtypes. Our results contribute to a better understanding of influenza tissue tropism across livestock species and its potential implications for zoonotic transmission.

**IMPORTANCE:** Since 2022, an unprecedented number of Influenza A virus (IAV) H5N1 detections have been reported in mammals. Sheep, goats, and alpacas are often raised in integrated production systems and may serve as hosts at the wildlife-livestock interface. The establishment of H5N1 in any mammalian host also increases its pandemic potential. Our results reveal species- and tissue-dependent patterns of H5N1 attachment and identify tissues that support viral binding. Furthermore, we show that respiratory tissues may be susceptible to other IAV subtypes, suggesting that these species could act as mixing vessels for reassortment events and the emergence of novel IAV strains. These results expand our understanding of the host range and tissue tropism of H5N1 and other IAV subtypes, providing a foundation for targeted surveillance of minor livestock species. Collectively, this work underscores the importance of considering diverse livestock hosts when evaluating the continued spread of H5N1 at the avian-mammalian interface.

## INTRODUCTION

Influenza A Viruses (IAVs) are negative-sense, single-stranded RNA viruses that belong to the Orthomyxoviridae family (1). Their genome has 8 segments that encode the major viral proteins, including the surface glycoproteins, Hemagglutinin (HA) and Neuraminidase (NA), which are also used to classify the virus into different subtypes (2). Because of the nature of IAV replication, the virus can rapidly mutate, leading to antigenic drift, in which minor point mutations can alter it over time. Furthermore, the segmented nature of the IAV genome facilitates antigenic shift, in which the virus can swap a gene segment for another following co-infection, as observed in previous pandemics (3, 4). Avian IAVs (AIVs) are also classified as low-pathogenic (LPAIV) or highly pathogenic (HPAIV) based on the presence of a polybasic cleavage site in the HA and the pathotype in chickens. The former allows cleavage by broad-host proteases and facilitates systemic infection in poultry (5, 6). HPAIVs are exclusively associated with the H5Nx or H7Nx subtype (7). Since 2022, HPAIV H5N1 subtype, clade 2.3.4.4b, has circulated continuously worldwide, including recent detections in Australia, marking the first incursion on that continent (8, 9). Strikingly, the current panzootic situation has marked a significant shift in the virus’s ecology, with more spillover events associated with H5N1 circulation documented across multiple mammalian species, including sea lions, rodents, cats, raccoons, and foxes (10, 11). The expanded host range of H5N1 is also well illustrated by spillover events into dairy cattle, which were confirmed in March 2024. As of today, more than 1,000 dairy herds across 20 states have been affected in the U.S (USDA-APHIS 2026), and serological evidence has been documented in the Netherlands (12–14). In addition, the 2024-2025 season saw record numbers of H5N1-associated human cases in the U.S., with most presenting with conjunctivitis and mild respiratory signs. This includes the first human fatality in the U.S. early in 2025 (15).

HPAIV H5N1 cases were also reported recently for the first time in goats and alpacas in the U.S. and sheep in the UK, reaffirming the expanded host range of H5N1 (16–18). In all cases, the animals were in close contact with poultry that had previously tested positive for HPAIV H5N1. For alpacas and goats, the disease was associated with increased mortality and abortions. Detection in sheep in the U.K. followed bulk milk tank testing of cattle and targeted surveillance of captive mammalian species on HPAIV-infected premises. Sheep, goats, and alpacas contribute to global agricultural production by providing meat, milk, fiber, and other animal-derived products (19). Therefore, the establishment of viral pathogens such as HPAIV H5N1 could have devastating consequences. Multiple cellular and viral factors drive the host range of influenza viruses (5). Among those, HA plays a major role in viral attachment and the beginning of the replication cycle in a susceptible cell by binding to sialic acid (SA) receptors on the cell surface (2, 20). Historically, influenza viruses of avian origin tend to bind to **α-**2,3-SAs, while mammalian influenza viruses favor binding to **α**-2,6-SAs. Importantly, the shift in these binding profiles has been determined to be a relevant feature in cross-species transmission at the avian-mammalian interface (2, 21). A well-characterized example is swine, where the dual presence of both ***α***-2,3-linked and **α**-2,6-linked sialic acid receptors facilitates infection by mammalian-and avian-derived viruses (22). This can potentially favor reassortment events and the emergence of novel IAV strains with unknown phenotypic features.

We previously examined the mammary glands of dairy cows from HPAIV H5N1 index cases to determine whether SA receptors were present in mammary tissue, potentially facilitating initial viral binding and entry into cells (21, 23). Furthermore, a follow-up study using mammary tissues from several livestock species, including pigs, beef cattle, dairy cattle, goats, alpacas and humans, also showed the presence of both SAs and efficient binding to multiple H5N1 strains of clade 2.3.4.4b (24). In this study, we sought to further characterize the SA distribution in recently affected livestock species, including sheep, goats, and alpacas, and to assess the binding profiles of H5N1 using a panel of formalin-fixed respiratory and non-respiratory tissues from each species. The results showed that a prototype clade 2.3.4.4b H5N1 strain efficiently binds to multiple tissues. We determined that binding efficacy is higher in sheep and goats than in alpacas. Interestingly, an ancestral strain from the Goose-Guangdong (Gs/Gd) lineage binds with efficiency comparable to that of our clade 2.3.4.4b prototype strain. Finally, the same experimental procedures were carried out with a panel of swine influenza viruses and a prototypical pandemic H1N1, showing that all strains bind to respiratory tissues, although differences in binding were observed.

## RESULTS

### Generation of recombinant low pathogenic H5N1 viruses

To explore the binding profiles of H5N1 across multiple tissues in sheep, goats, and alpacas, we used reverse genetics (RG) to generate recombinant LPAIV H5N1 viruses (Fig. 1A). To do so, an LPAIV version of a prototypical H5N1 virus from clade 2.3.4.4b (A/turkey/Minnesota/22-010654-002/2022 (H5N1)) and an LPAIV version of a prototype progenitor strain of the Goose-Guangdong (A/goose/Guangdong/1/1996) were each combined with their respective NA genes and the internal gene constellation of the laboratory-adapted strain A/Puerto Rico/08/34 (H1N1) (24). Both viruses (LP-gs/GD/96 (H5N1) and LP-tk/MN/22 (H5N1)) were successfully obtained by RG assays, and their growth dynamics were characterized in MDCK cells over 72 hours. The results (Fig. 1B) show that both strains reach high and comparable titers, with LP-gs/GD/96 (H5N1) at log_10_ 7.9 TCID50eq and LP-tk/MN/22 (H5N1) at log_10_ 7.7 TCID50eq by 72 hours. Statistically significant differences were observed only at 24 hours between the 2 viruses. Next, we investigated the SAs receptor-binding preferences of LP-gs/GD/96 (H5N1) and LP-tk/MN/22 (H5N1) using solid-phase binding assays with sialylated glycans representing α-2,3 and α-2,6 SAs receptors (Fig 1C-D). A/California/04/09 (H1N1)(pdm09) with known binding to α-2,6 SAs but not α-2,3 SAs was used as a control (25). In each case, a fixed amount of virus (128 HA units/50 μL) was incubated with increasing amounts of α-2,6 (Fig. 1C) and α-2,3 (Fig. 1D) SAs. The results show negligible binding of LP-gs/GD/96 (H5N1) and LP-tk/MN/22 (H5N1) to α-2,6 SAs, whereas pdm09 binds efficiently to these SAs. On the other hand, both H5N1 viruses evaluated bound efficiently to α-2,3 SAs, while marginal binding was observed from pdm09. Taken together, the results show the generation of LPAIV H5N1 and reaffirm the preference for α-2,3 SAs, consistent with previous reports (26, 27).

**Figure 1.**
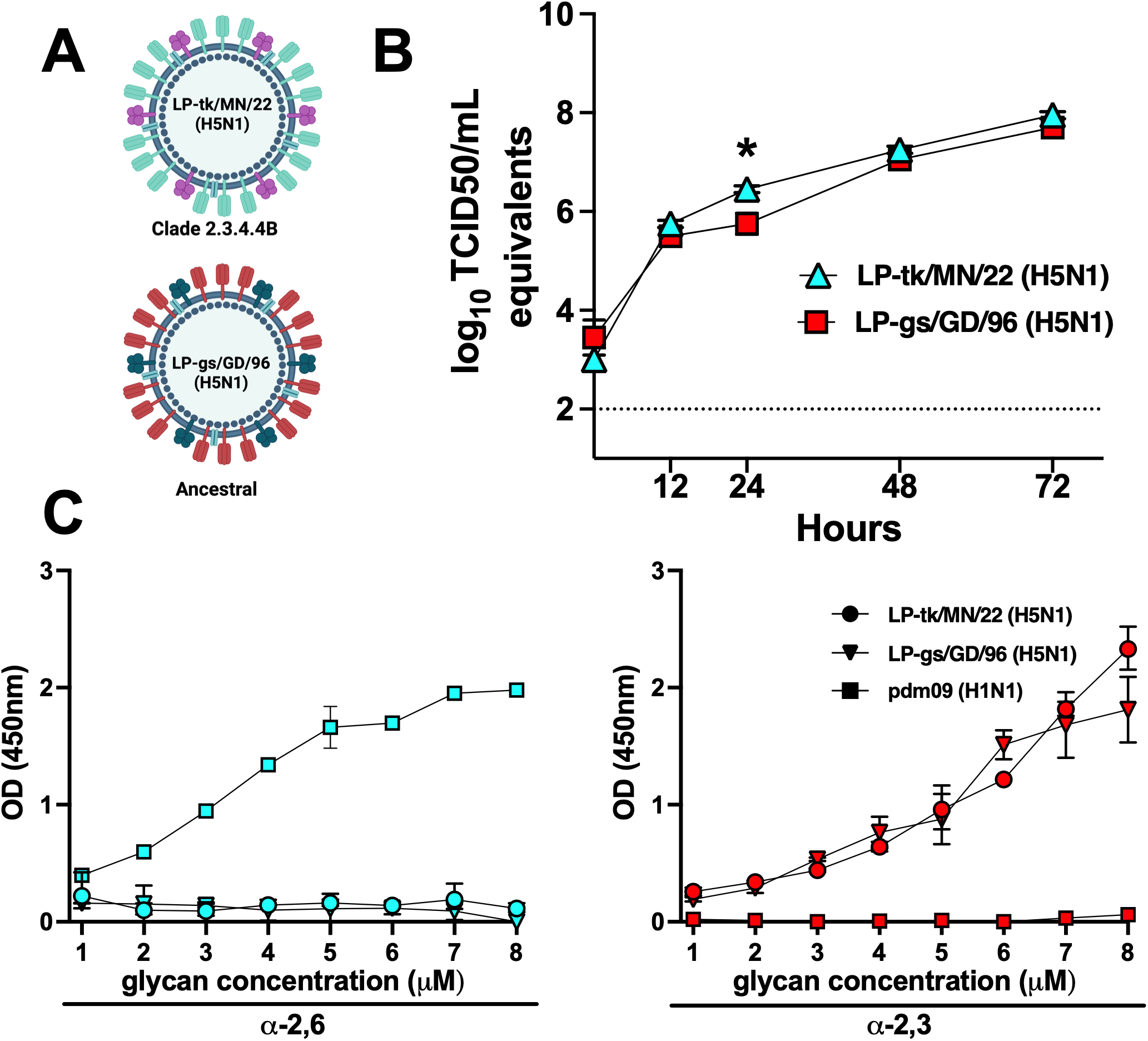
Characterization of the recombinant LPAIV H5N1 generated. **(A)** Schematic representation of the LPAIV viruses obtained by reverse genetics. The LPAIV versions of the HA of LP-tk/MN/22 (H5N1) clade 2.3.4.4b (top panel) and LP-gs/GD/96 (H5N1) (Gs/Gd lineage precursor; bottom panel) were combined with their respective NA genes and the internal genes of a PR8 (laboratory-adapted strain). **(B)** MDCK cells were inoculated with LP-tk/MN/22 (H5N1) (white triangles) or LP-gs/GD/96 (H5N1) (red squares) at an MOI of 0.01. Samples were collected at 0, 12, 24, 48, and 72 hours post-inoculation, and viral RNA levels were quantified by RT-qPCR and expressed as TCID50 equivalents using a standard curve of a virus with a known TCID50 titer. A p < 0.05 was considered significant. **(C)** LP-tk/MN/22 (H5N1) (circles), LP-gs/GD/96 (H5N1) (triangles) or pdm09 (H1N1) (squares) at a standard concentration of 128 HA units/50 μL were incubated with increasing concentrations of 6’ SLN-PAA-biot (left panel; white symbols) or 3’ SLN-PAA-biot (right panel; red symbols) in serial 2-fold dilutions (range: 10-0.078 μg/mL). The binding of each virus to the different concentrations was quantified by measuring the absorbance at 450nm.

### Distribution of influenza SA receptors in multiple tissues from sheep, goats and alpacas

To investigate the distribution of SA receptors (α-2,3 and α-2,6) in sheep, goats, and alpacas, we collected multiple tissues (conjunctiva, nasal turbinate, trachea, lung bifurcation, distal lung, esophagus, and rumen) from each species (n =3 per species) and processed them for routine histopathology evaluation. All the animals used were influenza-negative and overall healthy. Using these tissues, we investigated the presence of SA-α-2,3 and SA-α-2,6 using different lectins (Fig. 2). We employed a combination of Maackia amurensis I (magenta) and II (cyan) (MAL I &II) for the detection of SA-α-2,3, while Sambucus nigra (SNA; brown) was used for the detection of SA-α-2,6 (Table 1) (28). In all cases, hematoxylin and eosin slides were included in the analysis. The results show similar α-2,3 and α-2,6 SA distributions in the conjunctiva and nasal turbinates across the species evaluated (Table 2). A subset of differences was observed in the goat conjunctiva, with SNA present at higher levels than the combined presence of MAL I & II (mean scores: 4 for SNA, 1.5 for MAL I, and 2 for MAL II). In the alpaca’s conjunctiva, MAL II was significantly higher than MAL I (mean: 4 vs 0.67). Similarly, the alpaca’s nasal turbinate showed higher MAL II expression than MAL I. The analysis of tracheal tissues revealed a homogeneous distribution of the markers tested across species, with the presence of α-2,3 and α-2,6 SAs in the sections evaluated. Similar results were observed for the lung bifurcation and the bronchi. However, alpacas showed considerably higher MAL II presence than MAL I and SNA in the alpaca’s lung bifurcation (scores of 3.5, 2, and 1.5, respectively). A similar trend, with higher MAL II than MAL I, was observed in the alveoli of all species tested (scores of 4 vs 1 for sheep, 3.3 vs 1.7 for goat, and 4 vs 0 for alpaca). Similar levels of MAL II and SNA were observed in the alveoli of all tissues examined.

**Figure 2.**
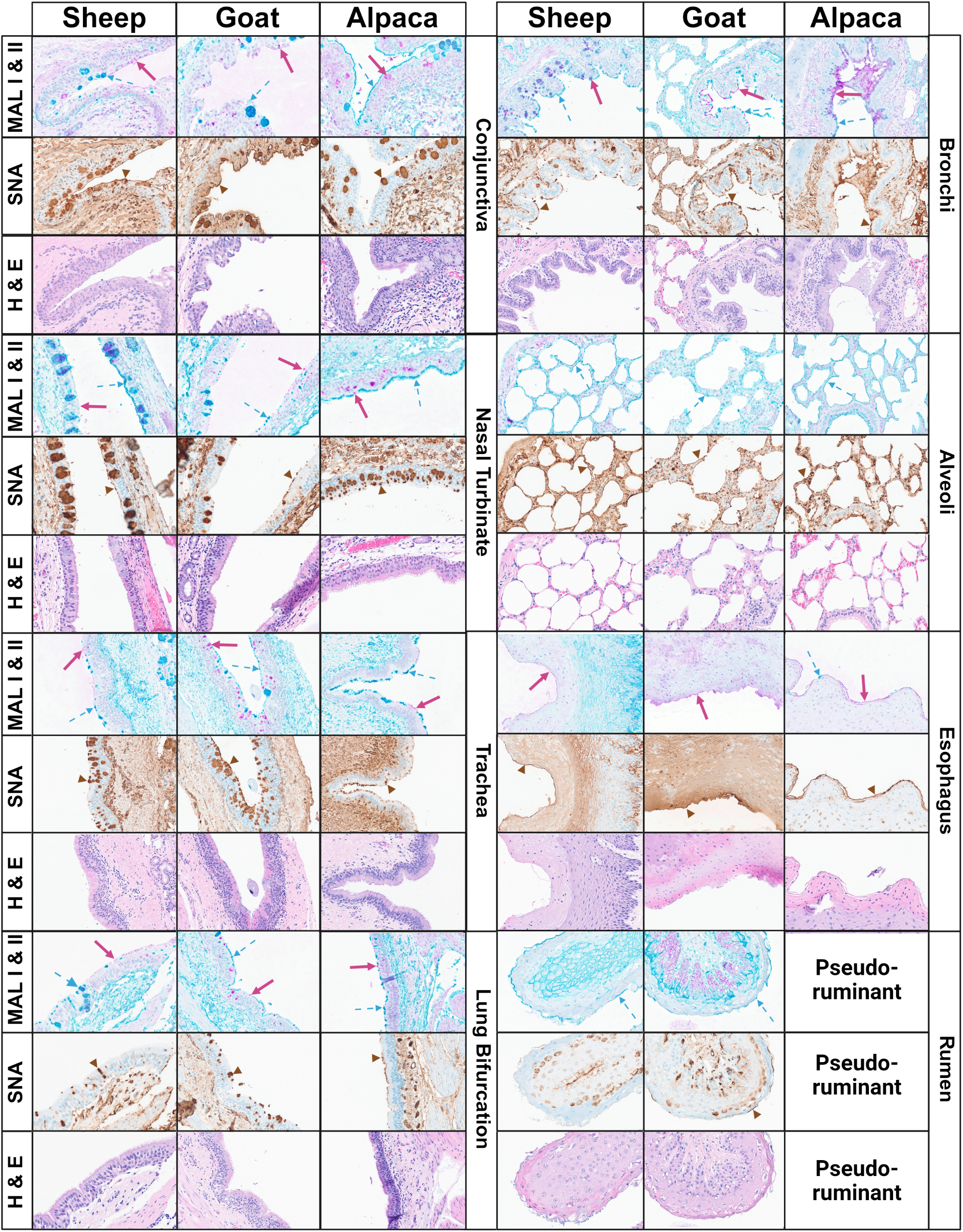
Sialic acid distribution in respiratory and non-respiratory tissues from sheep, goats and alpacas. The presence of α2,3- and α2,6-sialic acids (SAs) in conjunctiva, nasal turbinates, trachea, lung bifurcation, bronchi, alveoli, esophagus, and rumen was quantified using different lectins. MAL I and MAL-II were used to determine the presence of SA-α2,3, and SNA for SA-α2,6 (Table 1). In all cases, representative images of the analyzed independent tissues are shown. Arrows indicate regions of visible binding of the lectins to their respective receptors. Alpacas lack a rumen; therefore, no tissue was evaluated (29). At least 4 board-certified veterinary pathologists from the American College of Veterinary Pathologists evaluated all tissues, and results were reported using qualitative terms. Each receptor was assessed on a scale from 0 to 4 (Table 2).

**Table 1.** Lectin histochemistry and viral binding assay reagents.

| Item | Conjugate | Working concentration | Manufacturer | Product number |
| --- | --- | --- | --- | --- |
| Lectin staining |  |  |  |  |

| **Item** | **Conjugate** | **Working**<br>**concentration** | **Manufacturer** | **Product**<br>**number** |
| --- | --- | --- | --- | --- |
| *Maackia*<br>*amurensis*-I | Biotinylated | 20 µg/mL | Vector<br>Laboratories | B-1315-2 |
| *Maackia*<br>*amurensis*-II | Biotinylated | 4 µg/mL | Vector<br>Laboratories | B-1265-1 |
| *Sambucus nigra* | Biotinylated | 20 µg/mL | Vector<br>Laboratories | B-1305-2 |

**Table 2.** Summary of sialic acid lectin staining and LPAIV H5N1 binding in sheep, goat, and alpaca.

| *Animal* | *Binding* | *Conjunctiva* | *Localization* | *Mean* | *Binding* | *Nasal Turbinate* | *Localization* | *Mean* |
| --- | --- | --- | --- | --- | --- | --- | --- | --- |
| **Sheep** | α-2,3<br>MAL I | 4, -, - | E | 4 | α-2,3<br>MAL I | 2, 1, 2 | E | 1.67 |
| | α-2,3<br>MAL II | 2, -, - | GC | 2 | α-2,3<br>MAL II | 2, 3, 3 | E, GC | 2.67 |
| | α-2,6<br>SNA | 4, -, - | E, GC | 4 | α-2,6<br>SNA | 1, 3, 3 | GC | 2.33 |
|  | LP-tk/MN/22<br>(H5N1) | 3, -, - | E | 3 | LP-tk/MN/22<br>(H5N1) | 2, 2, 3 | E | 2.33 |
|  |  |  |  |  | LP-gs/GD/96<br>(H5N1) | 3, 4, 4 | SE | 3.67 |
| **Goat** | α-2,3<br>MAL I | -, 2, 1 | E, GC | 1.5 | α-2,3<br>MAL I | 1, 2, 1 | E | 1.33 |
| | α-2,3<br>MAL II | -, 2, 2 | E, GC | 2 | α-2,3<br>MAL II | 3, 2, 4 | E, GC | 3 |
| | α-2,6<br>SNA | -, 4, 4 | E, GC | 4 | α-2,6<br>SNA | 3, 3, 2 | GC | 2.67 |
|  | LP-tk/MN/22<br>(H5N1) | - ,1, 1 | E | 1 | LP-tk/MN/22<br>(H5N1) | 2, 1, 1 | E | 1.33 |
|  |  |  |  |  | LP-gs/GD/96<br>(H5N1) | 1, 1, 1 | E | 1 |
| **Alpaca** | α-2,3<br>MAL I | 1, 0, 1 | E, GC | 0.67 | α-2,3<br>MAL I | 1, 1, 0 | E | 0.67 |
| | α-2,3<br>MAL II | 4, 4, 4 | E, GC | 4 | α-2,3<br>MAL II | 3, 3, 4 | E, GC | 3.33 |

| | α-2,6<br>SNA | 4, 3, ,3 | E, GC | 3.33 | α-2,6<br>SNA | 2, 2, 3 | E, GC | 2.33 |
| --- | --- | --- | --- | --- | --- | --- | --- | --- |
|  | LP-tk/MN/22<br>(H5N1) | 3, 4, 3 | E | 3.33 | LP-tk/MN/22<br>(H5N1) | 3, 3, 3 | E | 3 |
|  |  |  |  |  | LP-gs/GD/96<br>(H5N1) | 4, 4, 4 | E | 4 |
| **Animal** | **Binding** | **Trachea** | **Localization** | **Mean** | **Binding** | **Lung**<br>**Bifurcation** | **Localization** | **Mean** |
| **Sheep** | α-2,3<br>MAL I | 1, 3, 2 | E | 2 | α-2,3<br>MAL I | 1, 2, 1 | E | 1.33 |
| | α-2,3<br>MAL II | 3, 2, 2 | E, GC | 2.33 | α-2,3<br>MAL II | 2, 2, 2 | E, GC | 2 |
| | α-2,6<br>SNA | 2, 4, 3 | E, GC | 3 | α-2,6<br>SNA | 1 ,2, 1 | GC | 1.33 |
|  | LP-tk/MN/22<br>(H5N1) | 3, 2, 2 | E | 2.33 | LP-tk/MN/22<br>(H5N1) | 4, 3, 3 | E | 3.33 |
|  | LP-gs/GD/96<br>(H5N1) | 2,1,2 | E | 1.67 |  |  |  |  |
| **Goat** | α-2,3<br>MAL I | 2, 2, 1 | E, GC | 1.67 | α-2,3<br>MAL I | 2, 2, 1 | E, GC | 1.67 |
| | α-2,3<br>MAL II | 3, 3, 4 | E, GC | 3.33 | α-2,3<br>MAL II | 3, 2, 3 | E, GC | 2.67 |
| | α-2,6<br>SNA | 4, 3, 3 | E, GC | 3.33 | α-2,6<br>SNA | 3, 3, 3 | E, GC | 3 |
|  | LP-tk/MN/22<br>(H5N1) | 3, 3, 2 | E | 2.67 | LP-tk/MN/22<br>(H5N1) | 3, 4, 3 | E | 3.33 |
|  | LP-gs/GD/96<br>(H5N1) | 1,2,1 | E | 1.33 |  |  |  |  |
| **Alpaca** | α-2,3<br>MAL I | 2, 2, - | E | 2 | α-2,3<br>MAL I | 2, -, 2 | E | 2 |
| | α-2,3<br>MAL II | 3, 3, - | E, GC | 3 | α-2,3<br>MAL II | 3, -, 4 | E | 3.5 |
| | α-2,6<br>SNA | 3, 2, - | E, GC | 2.5 | α-2,6<br>SNA | 1, -, 2 | E | 1.5 |
|  | LP-tk/MN/22<br>(H5N1) | 1, 2, - | E | 1.5 | LP-tk/MN/22<br>(H5N1) | 1, -, 1 | E | 1 |
|  | LP-gs/GD/96<br>(H5N1) | 0,1,- | E | 0.5 |  |  |  |  |
| **Animal** | **Binding** | **Bronchi** | **Localization** | **Mean** | **Binding** | **Alveoli** | **Localization** | **Mean** |
| **Sheep** | α-2,3<br>MAL I | 1, 2, 1 | E, GC | 1.33 | α-2,3<br>MAL I | 1,1,1 | E | 1 |
| | α-2,3<br>MAL II | 3, 3, 4 | E, GC | 3.33 | α-2,3<br>MAL II | 4,4,4 | E | 4 |
| | α-2,6<br>SNA | 3, 2, 3 | E, GC | 2.67 | α-2,6<br>SNA | 3,3,4 | E | 3.33 |
|  | LP-tk/MN/22<br>(H5N1) | 4, 4, 3 | E, GC | 3.67 | LP-tk/MN/22<br>(H5N1) | 4,4,4 | E | 4 |
|  | LP-gs/GD/96<br>(H5N1) | 4,4,3 | E,GC | 3.67 | LP-gs/GD/96<br>(H5N1) | 4,4,3 | E | 3.67 |
| **Goat** | α-2,3<br>MAL I | 3, ,3, 2 | E, GC | 2.67 | α-2,3<br>MAL I | 2, 1,2 | E | 1.67 |
| | α-2,3<br>MAL II | 3, 3, 4 | E, GC | 3.33 | α-2,3<br>MAL II | 4, 3, 3 | E | 3.33 |
| | α-2,6<br>SNA | 4, 3, 4 | E | 3.67 | α-2,6<br>SNA | 4,3,4 | E | 3.67 |
|  | LP-tk/MN/22<br>(H5N1) | 4,4,4 | E | 4 | LP-tk/MN/22<br>(H5N1) | 3,3,3 | E | 3 |
|  | LP-gs/GD/96<br>(H5N1) | 4,3,3 | E,GC | 3.33 | LP-gs/GD/96<br>(H5N1) | 3,3,3 | E | 3 |
| **Alpaca** | α-2,3<br>MAL I | 1,3,2 | E | 2 | α-2,3<br>MAL I | 0,0,0 | E | 0 |
| | α-2,3<br>MAL II | 4,3,4 | E | 3.67 | α-2,3<br>MAL II | 4,4,4 | E | 4 |
| | α-2,6<br>SNA | 1,2,3 | E | 2 | α-2,6<br>SNA | 3,4,3 | E | 3.33 |
|  | LP-tk/MN/22<br>(H5N1) | 1,2,2 | E | 1.67 | LP-tk/MN/22<br>(H5N1) | 3,2,3 | E | 2.67 |
|  | LP-gs/GD/96<br>(H5N1) | 2,1,1 | E,GC | 1.33 | LP-gs/GD/96<br>(H5N1) | 3,2,1 | E | 2 |
| **Animal** | **Binding** | **Esophagus** | **Localization** | **Mean** | **Binding** | **Rumen** | **Localization** | **Mean** |
| **Sheep** | α-2,3<br>MAL I | 2,1,1 | E | 1.33 | α-2,3<br>MAL I | 0,0,0 | E | 0 |
| | α-2,3<br>MAL II | 1,1,2 | E | 1.33 | α-2,3<br>MAL II | 3,3,3 | E | 3 |
| | α-2,6<br>SNA | 2,1,2 | E | 1.67 | α-2,6<br>SNA | 1,0,0 | E | 0.33 |
|  | LP-tk/MN/22<br>(H5N1) | 0,1,1 | E | 0.67 | LP-tk/MN/22<br>(H5N1) | 1,0,1 | E | 0.67 |
| **Goat** | α-2,3<br>MAL I | 2,2,2 | E | 2 | α-2,3<br>MAL I | 0,0,0 | E | 0 |
| | α-2,3<br>MAL II | 0,0,0 | E | 0 | α-2,3<br>MAL II | 3,2,2 | E | 2.33 |
| | α-2,6<br>SNA | 4,4,4 | E | 4 | α-2,6<br>SNA | 1,1,1 | E | 1 |
|  | LP-tk/MN/22<br>(H5N1) | 0,0,1 | E | 0.33 | LP-tk/MN/22<br>(H5N1) | 0,1,0 | E | 0.33 |
| **Alpaca** | α-2,3<br>MAL I | 2,1,2 | E | 1.67 | α-2,3<br>MAL I | n/a | n/a | - |
| | α-2,3<br>MAL II | 1,1,1 | E | 1 | α-2,3<br>MAL II | n/a | n/a | - |
| | α-2,6<br>SNA | 3,3,3 | E | 3 | α-2,6<br>SNA | n/a | n/a | - |
|  | LP-tk/MN/22<br>(H5N1) | 1,0,0 | E | 0.33 | LP-tk/MN/22<br>(H5N1) | n/a | n/a | - |

**E:** Tissue epithelial cells. **GC:** Goblet cells. **N/A:** Non-applicable (Pseudo-ruminant). **(-):** Not evaluated

We also investigated the α-2,3 and α-2,6 SAs distribution in non-respiratory tissues including esophagus and rumen (the latter only for sheep and goat since alpacas do not have a true rumen (29)). We chose these tissues since they are part of the digestive tract and ingestion of H5N1-contaminated elements could be an exposure pathway for these animals. Weak amounts of MAL II were seen in the goat esophagus, but MAL I, MAL II, and SNA were present in all other esophageal tissues examined. In the rumen of sheep and goats, negligible lectin staining was observed, except for MAL II in both species. Taken together, the results revealed the distribution of the α-2,3 and α-2,6 SAs across multiple tissues from sheep, goats, and alpacas. However, differences between MAL I and MAL II were evident in certain tissues, such as the alpaca conjunctiva, nasal turbinates, and bronchi, suggesting the presence of distinct α2,3 SA subpopulations.

### IAV H5N1 clade 2.3.4.4b binds to respiratory tissues of sheep, goats, and alpacas

We used LP-tk/MN/22 (H5N1) generated by RG to evaluate its binding profiles to sections of nasal turbinates, trachea, lung bifurcation, bronchi and alveoli of sheep, goats and alpacas (Fig. 3A). A standard concentration of LP-tk/MN/22 (H5N1) (100 HA units/50 μL) was incubated with the various tissues examined, and viral binding was assessed using an anti-HA antibody. In all cases, a control without virus was included to rule out potential non-specific binding. A 2^nd^ control treated with virus, but no primary antibody was also evaluated, showing lack of nonspecific binding of the secondary antibody (data not shown). The results revealed binding of LP-tk/MN/22 (H5N1) at various levels of the respiratory tract, with moderate to strong binding noted at the upper respiratory mucosa lining the nasal turbinates, mild to moderate binding in the tracheal mucosa and bronchial mucosa, and strong binding at the lower respiratory tissues of the bronchioles and alveoli (Fig. 3A, brown arrows). Interestingly, stronger binding of LP-tk/MN/22 (H5N1) was observed in the nasal turbinates of alpacas compared to sheep and goats. In contrast, viral binding to the trachea was multifocal and moderate in sheep and goats but weak in alpaca tissues. At the level of the lung bifurcation, diffusely strong-to-very-strong binding was observed in sheep and goats. However, alpaca showed multifocal, weak-to-moderate binding. The bronchi showed similar patterns as at the lung bifurcation. Finally, viral binding was very strong and diffuse in sheep alveoli (score of 4 out of 4). Overall, binding to the alveoli was lower in goat and alpaca compared to sheep.

**Figure 3.**
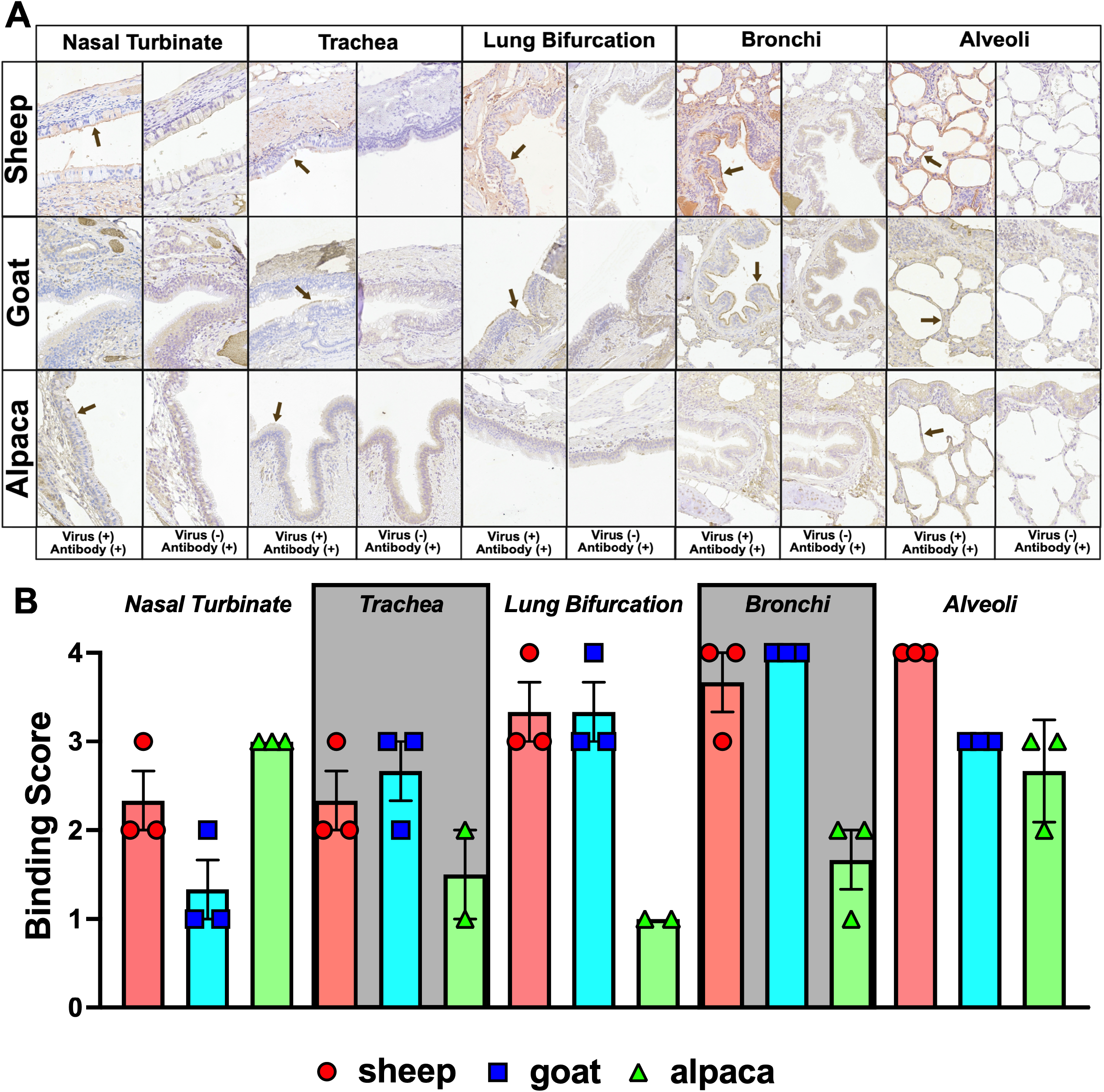
Binding of LP-tk/MN/22 (H5N1) to respiratory tissues of sheep, goats, and alpaca. **(A)** Representative pictures of virus binding using LP-tk/MN/22 (H5N1) clade 2.3.4.4b to nasal turbinates, trachea, lung bifurcation, bronchi, and alveoli. In all cases, a standard concentration of virus (100 HA units/50 μL) was used. Viral binding was visualized using a monoclonal anti-HA antibody (Clone CR9114). Control experiments were conducted in the absence of the virus to rule out nonspecific binding. Arrows indicate regions of representative virus-binding on the different tissues. At least 4 board-certified veterinary pathologists from the American College of Veterinary Pathologists evaluated all tissues, and results were reported using qualitative terms. Virus-binding was assessed on a scale from 0 to 4 (Table 2). <u>(Areas of strong viral binding are indicated by a</u> <u>brown arrow).</u> **(B)** Graph showing the binding scores for the independent tissues evaluated in each case from sheep (red), goat (blue), and alpacas (green).

The summary of the scores (Fig. 3B) for each evaluated section (n=3/tissue) is consistent with the representative images shown, in which higher binding to the alpacas’ nasal turbinate (green bars) was observed than in sheep and goats (red and blue bars, respectively). After that, there is a change in binding patterns, with higher binding of LP-tk/MN/22 (H5N1) to the trachea, lung hilum, secondary bronchi, and alveoli in sheep and goats compared to alpaca. Alveoli show more similar virus binding across the 3 species. Virus binding patterns match the distribution of sialic acid receptors. These findings may support efficient viral attachment across multiple ruminant species, suggesting susceptibility of both upper and lower respiratory tissues to infection, with pronounced binding to the lower airways and lungs, which may facilitate respiratory disease and viral replication following exposure.

### IAV H5N1 clade 2.3.4.4b binds to non-respiratory tissues of sheep, goats and alpacas with different efficiencies

After we established binding profiles for the respiratory tissues, we examined viral binding to the conjunctiva, esophagus, and rumen in each species, all of which are considered non-respiratory tissues (Fig. 4A). An important feature of HPAIVs is their capacity to infect non-respiratory tissues and cause systemic infections (3). Therefore, understanding the binding profile in non-respiratory tissues is important for assessing overall susceptibility to H5N1. The standard concentration of LP-tk/MN/22 (H5N1) (100 HA units/50 μL) was incubated with the various tissues examined, and viral binding was assessed using an anti-HA antibody. In all cases, a control without virus was included to rule out potential nonspecific binding. A 2^nd^ control with virus but no primary antibody was also evaluated, showing lack of nonspecific binding of the secondary antibody (data not shown). Viral binding to the LP-tk/MN/22 (H5N1) strain in the conjunctival mucosa was diffusely strong in sheep and alpaca, in sharp contrast to goats, where binding was scattered and weak. Next, we examined LP-tk/MN/22 (H5N1) viral binding across the gastrointestinal tract, revealing weaker binding than in the respiratory tract. The esophageal mucosa showed marginal binding across all species evaluated, with scattered, weak binding in sheep and scattered, weak-to-no binding in goats and alpacas. Virus binding in the rumen revealed similar findings, with weak to no binding in sheep and goats. It should be noted that alpacas are camelid pseudo-ruminants and lack a true rumen (29). The analysis of the numerical scores (Fig. 4B) shows high and similar LP-tk/MN/22 (H5N1) binding to the conjunctiva tissue of sheep (red bars) and alpaca (green bars), while weak binding is observed for goat (blue bars).

**Figure 4.**
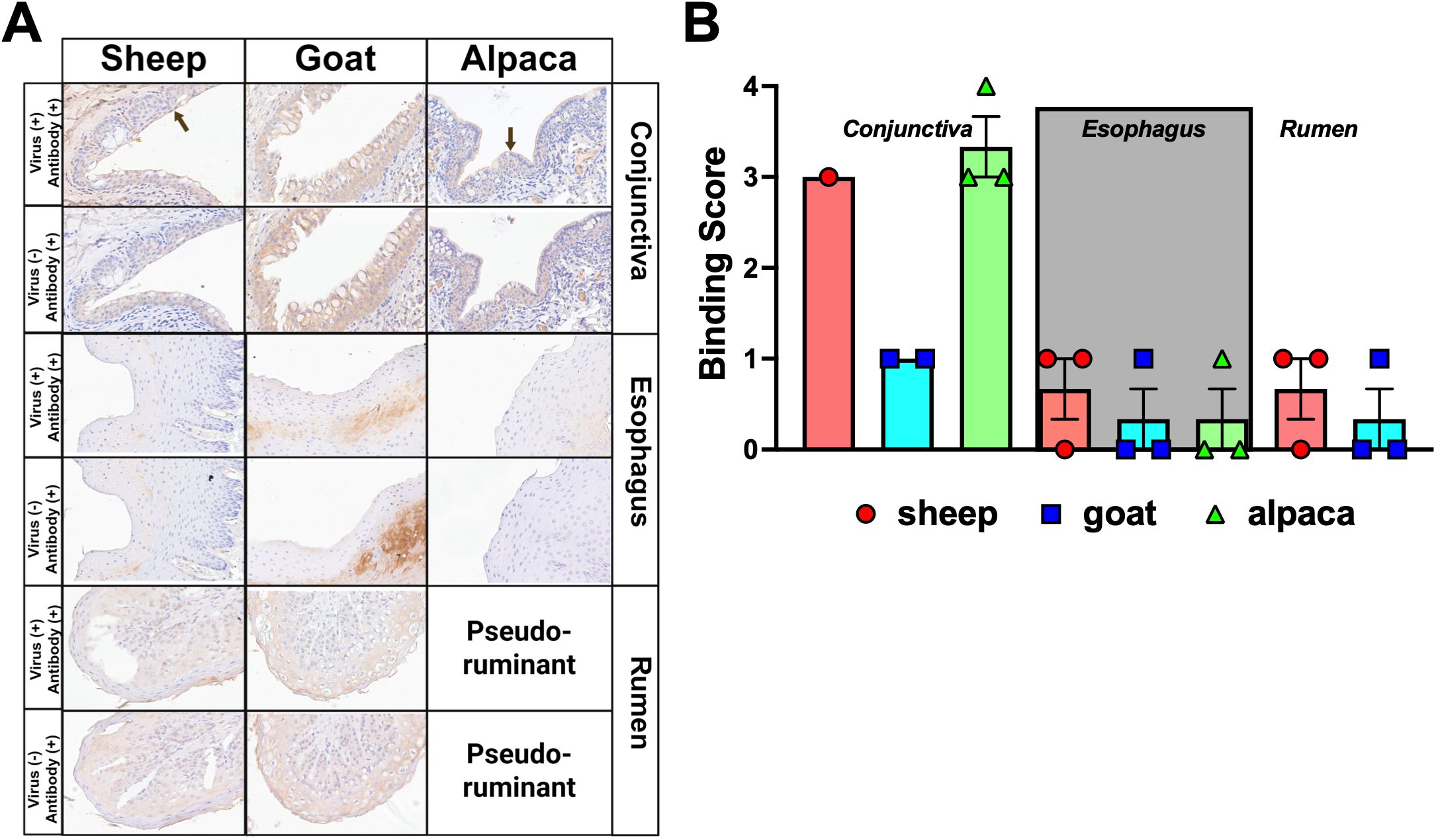
Binding of LP-tk/MN/22 (H5N1) to non-respiratory tissues of sheep, goats, and alpaca. **(A)** Representative pictures of virus binding using LP-tk/MN/22 (H5N1) clade 2.3.4.4b to conjunctiva, esophagus, and rumen. The latter was only analyzed for sheep and goats since alpacas do not have a rumen (29). In all cases, a standard concentration of virus (100 HA units/50 μL) was used. Viral binding was visualized using a monoclonal anti-HA antibody (Clone CR9114). Control experiments were conducted in the absence of the virus to rule out nonspecific binding. Arrows indicate regions of representative virus-binding on the different tissues. At least 4 board-certified veterinary pathologists from the American College of Veterinary Pathologists evaluated all tissues, and results were reported using qualitative terms. Virus-binding was assessed on a scale from 0 to 4 (Table 2). **(B)** Graph showing the binding scores for the independent tissues evaluated in each case from sheep (red), goat (blue), and alpacas (green).

For esophagus and rumen, a marginal score (score <1) was recorded for LP-tk/MN/22 binding regardless of the species evaluated. Collectively, these findings suggest that the conjunctiva is a viable route of entry and potential site of replication for H5N1, particularly in sheep and alpacas, but less so in goats. Furthermore, the gastrointestinal tract appears to play a limited role in viral tropism and susceptibility to H5N1.

### A prototype progenitor of the H5N1 Gs/Gd lineage efficiently binds to the respiratory tissues of sheep, goats, and alpacas

Since 2022, H5N1 infections have shown an expanded host range (30–32). A hypothesis to explain this expanded host range relates to the HA protein and potential features in the HA of circulating strains that enable binding to an expanded set of tissues from different animals, which were not present in previous circulating strains (33). To test this hypothesis in our system and evaluate potential differences at the level of viral binding, we interrogated different respiratory tissues from sheep, goat, and alpacas for the binding of LP-gs/GD/96 (H5N1), an LPAIV version of the prototype root progenitor of the Gs/Gd lineage (A/goose/Guangdong/1/1996) (34) (Fig. 5A). As in the experiments with LP-tk/MN/22 (H5N1) (Fig. 3 and 4), a standard concentration (100 HA units/50 μL) of virus was inoculated into different tissues, and the presence of viral binding was analyzed using an anti-HA antibody. In all cases, a control without virus was included to rule out potential nonspecific binding. A 2^nd^ control with virus but no primary antibody was also evaluated, showing lack of nonspecific binding of the secondary antibody (data not shown). The results show binding patterns like those previously described with LP-tk/MN/22 (H5N1) (Fig. 3 and 4). Viral binding in the nasal turbinates was diffuse and very strong in sheep and alpacas. However, scattered, weak binding was observed in nasal turbinates from goats. In the trachea, LP-gs/GD/96 (H5N1) showed moderate-to-weak multifocal binding in sheep and goats, with marginal binding in alpaca tracheal tissues. For the bronchi, overall binding was higher than in the trachea. However, the binding levels did not reach those observed in the nasal turbinates. Viral binding on bronchi was diffusely strong to very strong for sheep, diffusely strong for goats, and multifocal and moderate for alpaca. Finally, LP-gs/GD/96 (H5N1) binding to alveolar epithelium was diffuse and very strong in sheep, diffuse and strong in goats, and diffuse and weak in alpaca.

**Figure 5.**
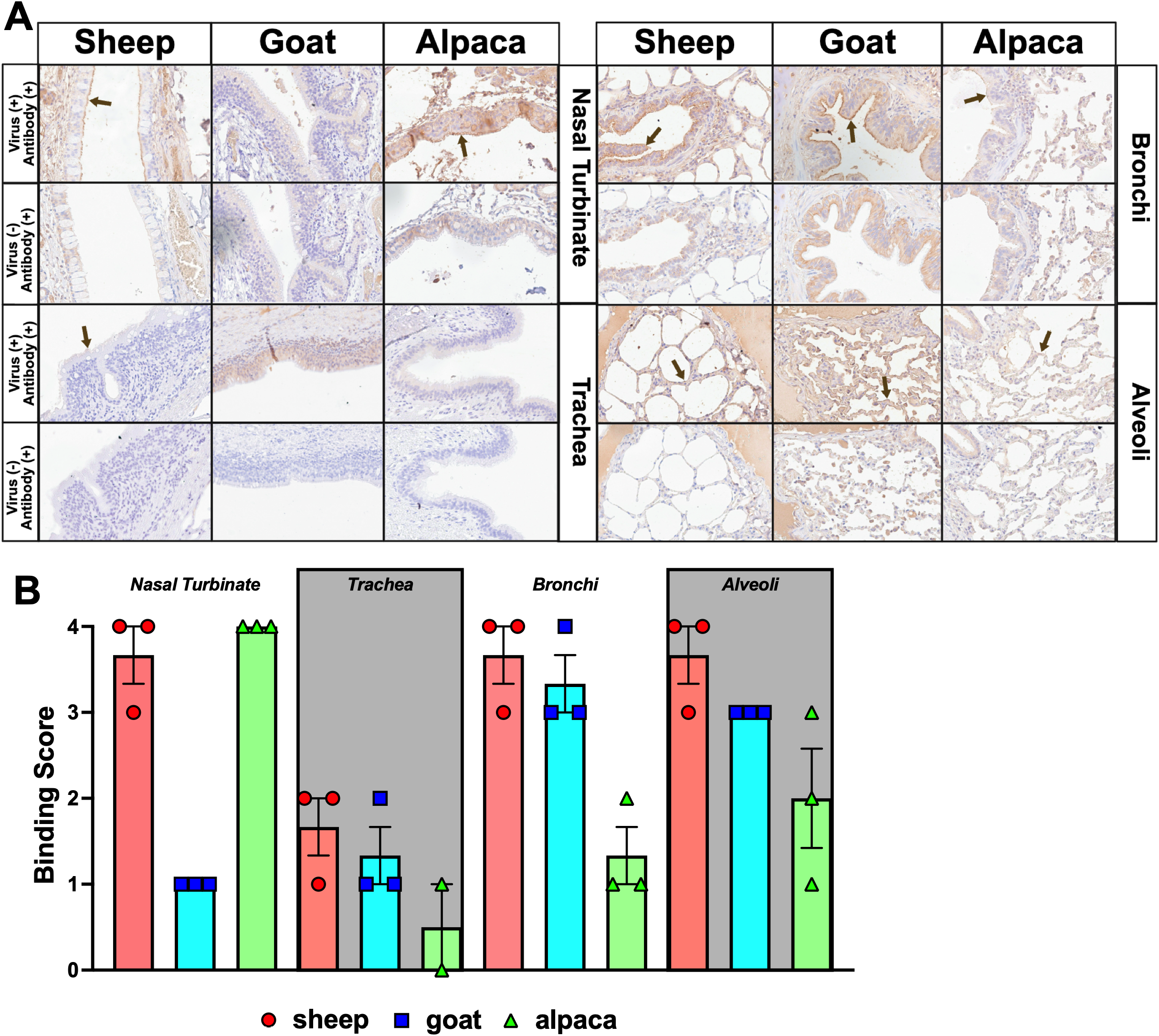
Binding of LP-gs/GD/96 (H5N1) to respiratory tissues of sheep, goats and alpacas. **(A)** Representative pictures of virus binding using LP-gs/GD/96 (H5N1) (Gs/Gd lineage precursor) to nasal turbinates, trachea, bronchi and alveoli. In all cases, a standard concentration of virus (100 HA units/50 μL) was used. Viral binding was visualized using a monoclonal anti-HA antibody (Clone CR9114). Control experiments were conducted in the absence of the virus to rule out nonspecific binding. Arrows indicate regions of representative virus-binding on the different tissues. At least 4 board-certified veterinary pathologists from the American College of Veterinary Pathologists evaluated all tissues, and results were reported using qualitative terms. Virus-binding was assessed on a scale from 0 to 4 (Table 2). **(B)** Graph showing the binding scores for the independent tissues evaluated in each case from sheep (red), goat (blue), and alpacas (green).

The summary of the binding scores (Fig. 5B) confirms the patterns observed with strong binding of LP-gs/GD/96 (H5N1) to the nasal turbinates of sheep and alpacas but not goats, weak to moderate binding in tracheal tissues across all the species evaluated, strong binding to sheep’s and goat’s bronchi with a decline to moderate for alpacas, and strong to very strong binding to alveolar tissues of sheep and goats but only moderate binding in the context of alpacas. Taken together, the results demonstrate that LP-gs/GD/96 (H5N1) exhibits highly similar patterns of tissue binding across sheep, goats, and alpacas compared to LP-tk/MN/22 (H5N1), characterized by strong viral attachment throughout the respiratory tract and particularly robust binding in the bronchi and alveoli for sheep and goats. These findings suggest that receptor distribution and tissue tropism supporting H5N1 attachment have been conserved across Gs/Gd strains despite viral evolution, indicating a consistent capacity to infect the lower respiratory tract in these species.

### Different IAV subtypes bind efficiently to respiratory sections of sheep and goats

The SA distribution (Fig. 2) and IAV H5N1 binding to the tissues of the evaluated species (Figs. 3-5) suggest that other IAV subtypes could bind similarly and potentially replicate. In this scenario, co-infection with H5N1 and more mammalian-adapted IAV subtypes could be particularly concerning due to the risk of reassortment. A classic example is swine, often called “mixing vessels” because they are susceptible to both avian and mammalian IAV strains. This provides an environment for reassortment events and the emergence of novel IAV strains with unknown phenotypic features, as reported for the emergence of the latest IAV pandemic in 2009 (35, 36). Therefore, we sought to investigate the binding capacity of IAV strains beyond H5N1 using bronchi (representative of the lower respiratory tract) from sheep and goats. Alpaca’s tissues were not included in the analysis due to the overall lower binding of H5N1 observed (Fig. 3 and 5). We performed viral binding assays with LP-tk/MN/22 (H5N1) and LP-gs/GD/96 (H5N1) together with pdm09 (H1N1), a prototypical pandemic H1N1 strain derived from humans. Furthermore, we included two different swine influenza strains: A/swine/Minnesota/A02245397/2020 (H1N1) (swH1N1) and A/swine/Iowa/A022861497/2023 (H3N2)(swH3N2). Pdm09 (H1N1), swH1N1, and swH3N2 were generated by RG, and a standard concentration (100 HA units/50 μL) of each one of those, together with LP-tk/MN/22 (H5N1) and LP-gs/GD/96 (H5N1) virus, was incubated with bronchi sections of sheep and goat. The presence of viral binding was analyzed using an anti-HA antibody. In all cases, a control without virus was included to rule out potential nonspecific binding. A 2^nd^ control with virus but no primary antibody was also evaluated, showing lack of nonspecific binding of the secondary antibody (data not shown).

The results (Fig. 6A) showed that, consistent with previous results, LP-tk/MN/22 (H5N1) showed diffusely strong-to-very-strong binding, and LP-gs/GD/96 (H5N1) showed a similar pattern. For pdm09 (H1N1), a pattern like that observed for LP-tk/MN/22 (H5N1) was observed, with strong to very strong binding to bronchial tissues from sheep and goats, respectively. More interesting were the results for the swine IAV strains, where binding was consistently very strong in both cases, with a trend toward higher binding than even the H5N1 strains examined. The summary of the scores (Fig. 6B) showed, as previously determined, binding for LP-tk/MN/22 (H5N1) and LP-gs/GD/96 (H5N1) between 3 (strong) and 4 (very strong), with similar patterns for pdm09 (H1N1). For swH1N1 and swH3N2, all examined tissues showed the maximum binding score (4 out of 4), indicating even stronger binding than that observed with the H5N1 strains. These findings indicate that the lower respiratory tract of small ruminants contains abundant receptors that support attachment by both avian and mammalian influenza viruses, suggesting a conserved susceptibility from the binding perspective to IAV infection.

**Figure 6.**
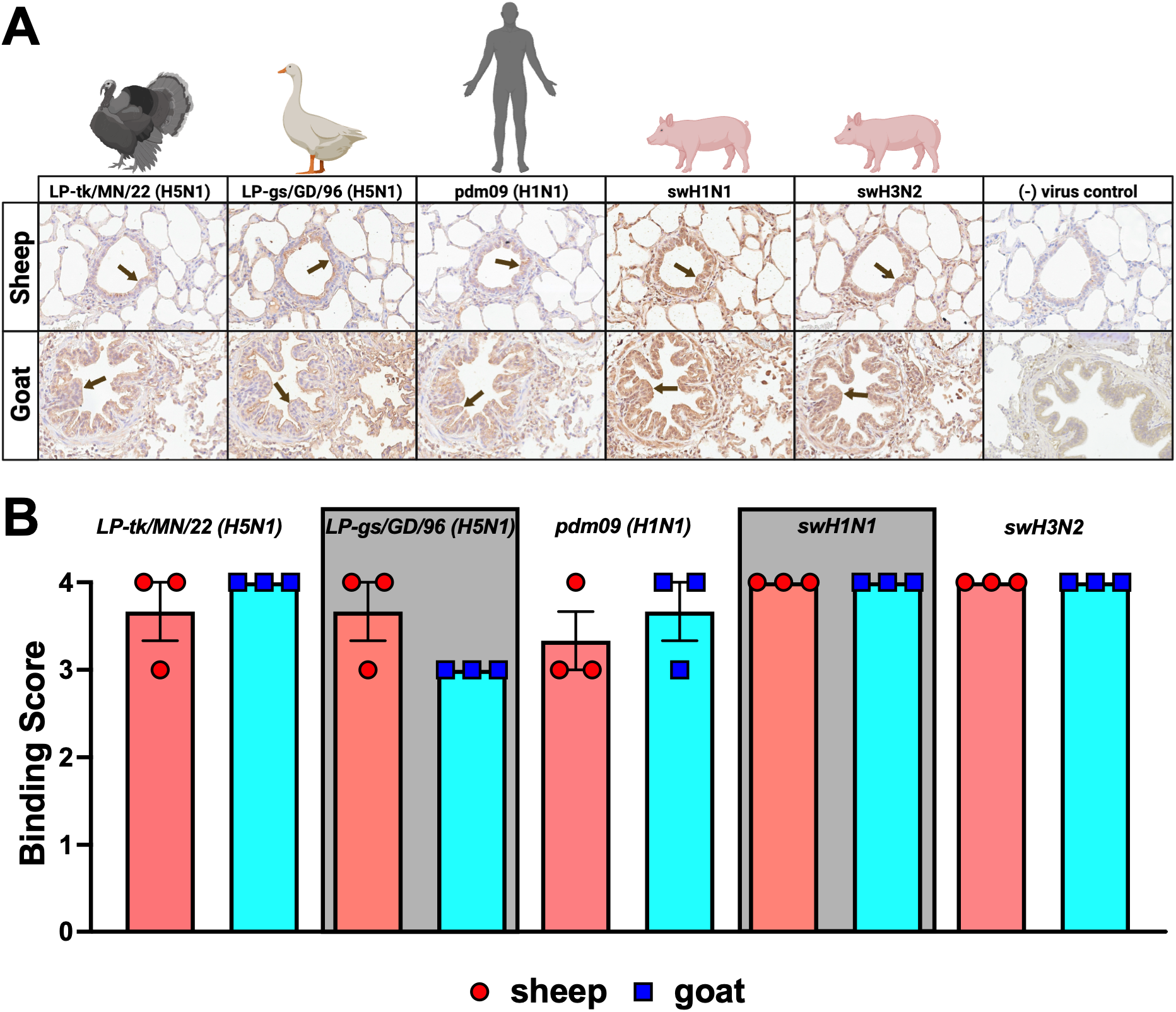
Viral binding of different IAVs to alveoli of sheep and goats. **(A)** Representative pictures of virus binding using LP-tk/MN/22 (H5N1), LP-gs/GD/96 (H5N1), pdm09 (H1N1), swH1N1, and swH3N2 viruses to alveoli sections of sheep and goats. In all cases, a standard concentration of virus (100 HA units/50 μL) was used. Viral binding was visualized using a monoclonal anti-HA antibody (Clone CR9114). Control experiments were conducted in the absence of the virus to rule out nonspecific binding. Arrows indicate regions of representative virus-binding on the different tissues. At least 4 board-certified veterinary pathologists from the American College of Veterinary Pathologists evaluated all tissues, and results were reported using qualitative terms. Virus-binding was assessed on a scale from 0 to 4 (Table 3). **(B)** Graph showing the binding scores for the independent tissues evaluated in each case from sheep (red) and goat (blue).

**Table 3.** Summary of IAV binding of multiple subtypes to bronchi tissues of sheep and goats.

| *Animal* | *Binding* | *Bronchi* | *Localization* | *Mean* |
| --- | --- | --- | --- | --- |
| **Sheep** | LP-tk/MN/22 (H5N1) | 4,4,3 | E | 3.67 |
|  | LP-gs/GD/96 (H5N1) | 4,4,3 | E | 3.67 |
|  | pdm09 (H1N1) | 3,4,3 | E | 3.33 |
|  | swH1N1 | 4,4,4 | E | 4 |
|  | swH3N2 | 4,4,4 | E | 4 |
| **Goat** | LP-tk/MN/22 (H5N1) | 4,4,4 | E | 4 |
|  | LP-gs/GD/96 (H5N1) | 3,3,3 | E | 3 |
|  | pdm09 (H1N1) | 4,4,3 | E | 3.67 |
|  | swH1N1 | 4,4,4 | E | 4 |
|  | swH3N2 | 4,4,4 | E | 4 |

## DISCUSSION

Since 2022, more than 200 million commercial birds have been affected by HPAIV H5N1 in the U.S. alone, significantly impacting the poultry industry and resulting in millions of dollars in economic losses and disruption of the food supply. Furthermore, an expanded host range, particularly in mammalian species, has been observed recently, with several species that were previously unaffected or only slightly affected now showing detection and, in some cases, severe disease, including death (11, 30, 37). This is well exemplified by the first detection of HPAIV H5N1 in dairy cows in 2024, marking a clear expansion of the viral host range and the first mammalian species in which H5N1 has established itself, with active transmission among animals (21, 24, 38). More importantly, the pathogenesis of H5N1 in dairy cows shows a shift in tissue tropism, with a change in the classical disease presentation of the virus toward a much more pronounced compromise of the mammary glands, characterized by reduced milk production, high viral titers in milk and mastitis rather than a predominant respiratory disease component (39–41). Because of this, several groups examined the effectiveness of pasteurization procedures in inactivating IAV, showing that, despite high viral loads, the process is effective (12, 42). This highlights the consequences at multiple levels of the expanded host range of H5N1 in areas such as animal and public health. HPAIV H5N1 cases were reported for the first time in sheep, goats, and alpacas (16–18). These species contribute to global agricultural production by providing meat, milk, fiber, and other animal-derived products (19). Therefore, the establishment of viral pathogens such as HPAIV H5N1 could have devastating consequences. Using virus-binding assays on a panel of respiratory and non-respiratory tissues from these species, we showed that a prototypical H5N1 clade 2.3.4.4b virus binds to different tissues across the different species evaluated (Fig. 7). Sheep and alpacas showed strong viral binding to tissues associated with the upper respiratory tract (nasal turbinates), whereas goats showed limited viral binding. The conjunctiva (another potential exposure route for the virus) showed consistently high viral binding for sheep and alpacas, with a sharp decrease for goats. In the lower respiratory tract, sheep and goats showed robust viral binding at the lung bifurcation and within the lungs (bronchi + alveoli), whereas lower viral binding was observed in tissues from alpacas. Negligible binding was observed in digestive tract tissues such as the esophagus and rumen.

**Figure 7.**
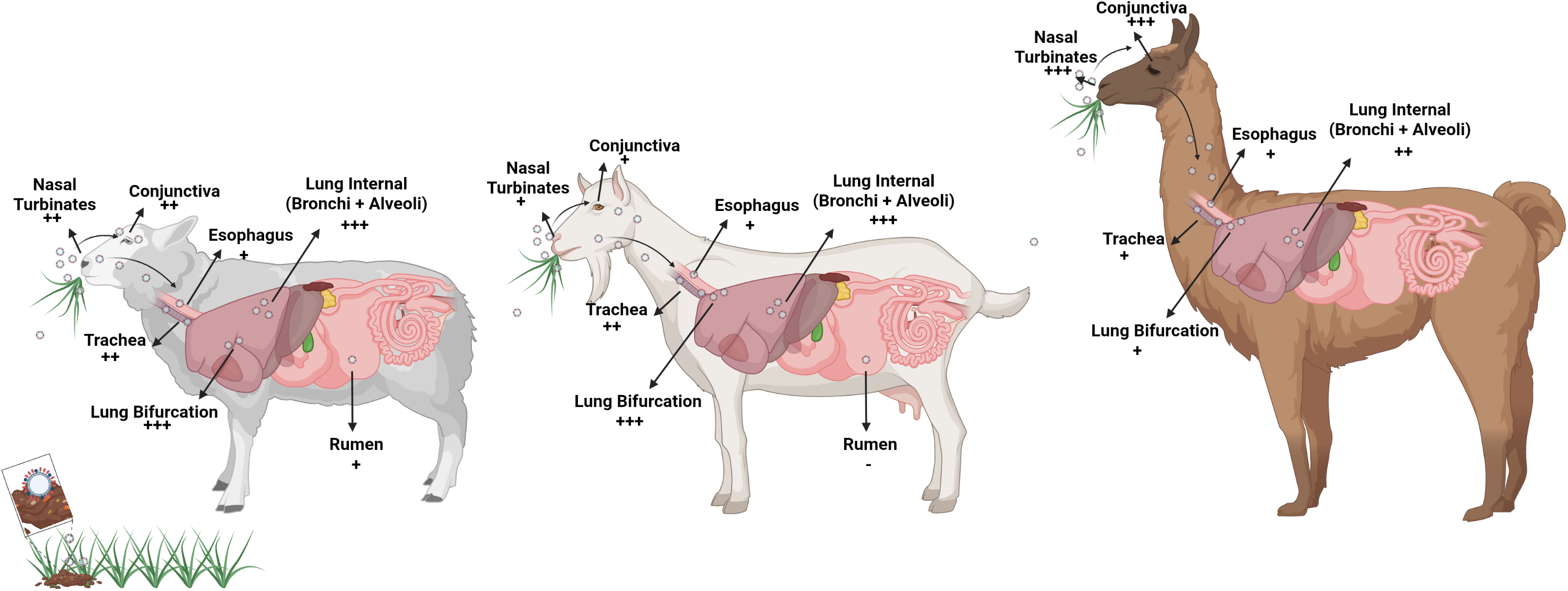
Summary of the virus-binding results obtained with LP-tk/MN/22 (H5N1). Schematic representation of sheep (left), goat (middle), and alpaca (right) showing the binding intensities (-, +, ++, +++) of LP-tk/MN/22 (H5N1) clade 2.3.4.4b to all the tissues evaluated in this report.

Many farms in the US practice mixed animal rearing, and the potential entry of HPAIV H5N1 onto farms may enable interspecies transmission, as evidenced in Oregon, where after a poultry farm tested positive for HPAIV, subsequent testing confirmed that pigs on the premises were also positive for H5N1. This marked the first time that HPAIV H5N1 was detected in swine in the US (43). The proximity of multiple susceptible species in agricultural settings can increase opportunities for repeated exposure and interspecies transmission, as these animals often share pasture, housing, and other resources. The susceptibility of livestock species, such as those investigated here, which have historically received less attention in influenza surveillance than poultry or waterfowl, is increasingly important as the host range of contemporary H5N1 viruses expands (44). Previous studies have also shown that stability significantly decreases after pasteurization in the products derived from H5N1-positive animals. However, in raw products, the virus can remain viable for extended periods (12, 45). This highlights the need to monitor infections and better understand H5N1 replication in these minor livestock species, whose milk products are often consumed raw, posing a particular concern for H5N1 introduction into the food production system, which is significantly less controlled than bovine dairy production (46).

The replication and transmission of HPAIV H5N1 across mammalian species are of great concern due to the potential for the virus to adapt to mammalian environments through genomic changes that could enhance replication and transmission, thereby increasing H5N1’s pandemic potential (47). Multiple molecular features, particularly those associated with the HA and the viral polymerase complex (PB2, PB1, and PA), have been shown to positively modulate H5N1 pathogenesis in mammalian systems (5, 48). Binding the virus to the host cell is a critical step in which avian-origin IAVs preferentially bind to α-2,3 SAs while mammalian-adapted IAVs bind more efficiently to α-2,6 SAs. Therefore, the shift in SA preference has been proposed as a critical feature in the host range and H5N1 capacity to infect mammalian hosts (21, 49). Key amino acids within the HA have been suggested as critical in the SA preference, including position 160, 226, and 228 (H3 numbering) (50–52). Among those, position 226 and the glutamine-to-leucine substitution (Q226L) are the most relevant changes reported to be associated with SA preference shifts, including in recent isolates (51, 53). Despite this, our results show that LP-tk/MN/22 (H5N1) and LP-gs/GD/96 (H5N1) retain the α-2,3 SAs with negligible binding to α-2,6 SAs (Fig. 1), consistent with other reports (26, 54). Furthermore, our viral-binding assays and the comparison between the binding profiles LP-tk/MN/22 (H5N1) (clade 2.3.4.4b) and LP-gs/GD/96 (H5N1)(Gs/Gd lineage precursor) (Fig. 3 and 5) suggest that the expanded host range observed in the last few years associated with H5N1 cannot be explained by the ability of the HA to bind to different tissues. This also suggests that the polymerase complex may be a more critical determinant of sustained viral replication in mammals and, potentially, of the expanded host range observed. Different genomic changes, particularly in PB2, have been associated with this, including Q to arginine at position 591 (Q591R), glutamic acid to lysine at position 627 (E627K), and aspartate to asparagine at position 701 (D701N) (55, 56). This is well illustrated by the recent H5N1 outbreak in dairy cows, where E627K is initially absent from bovine sequences. However, they emerged in human infections or in subsequent cattle-related surveillance (57). Like this, Q591R and D701N have been suggested as adaptation markers in avian-to-cattle transmission (58). Taken together, our results are consistent with the idea that early steps in the viral replicative cycle, such as binding to host cells, are not the primary features that enable circulating H5N1 strains to infect multiple species, as previously reported.

The results suggest that not only can H5N1 viruses bind to the respiratory tissues of sheep and goats, but other IAV subtypes (pdmH1N1, swH1N1 and swH3N2) have the same capacity (Fig. 6). Little to no information is available about the capacity of IAVs of the H1N1 or H3N2 subtype to infect sheep or goats. However, serological studies have shown 8% seropositivity to H1N1 in sheep but none in goats (59). Still, infections in sheep and goats are considered rare and probably usually overlooked. The recent emergence of H5N1 infections and the evident ability of more mammalian-adapted strains to bind host cells in these species suggest that this should be revisited (16–18). Minor livestock species, such as those evaluated here, could serve as mixing vessels, as has been extensively described for swine in the IAV ecology (60, 61). In particular, the replication of swine influenza in sheep and goats will be pursued further in experiments beyond the scope of the present report, given the strong binding observed in all animals examined. Recently, it was shown that swine influenza can also infect Holstein calves (62). This reinforces the idea of oversight across multiple species within the full spectrum of IAV’s host range.

Our results provide proof of principle that viral binding to the host cell is not a limitation on the virus’s eventual invasion of the host cell in the respiratory tract and other tissues. Still, the current report has inherent limitations that should be recognized. No active replication was evaluated in this report using the panel of tissues available to us. However, several groups have reported the experimental infection of sheep, goats, and alpacas with HPAIV H5N1. Intramammary inoculation in lactating sheep with H5Nx clade 2.3.4.4b (H5N1 or H5N5) caused clinical mastitis, high viral loads in milk, and transmission to suckling lambs. Moreover, aerosol exposure of nonlactating sheep led to transient respiratory infection, with low-level viral replication and seroconversion (63). In goats, H5N1 infection with different genotypes (B3.13 and B1.2) resulted in severe clinical mastitis and viral shedding in milk, with subsequent transmission to suckling kids. Viral RNA was detected in nasal and oral swabs and various tissues, and virus-neutralizing antibodies were present in serum, milk, and bronchoalveolar lavage fluid (18). Experimental infections have also been recently reported in alpacas, with an overall mild disease course and a slight increase in body temperature. Interestingly, the virus was only detected in the upper respiratory tract (64). These observations are consistent with our virus-binding results, in which binding to alpaca tissues is generally lower than to sheep and goats. However, strong binding is observed in the upper respiratory tract (Fig. 3). Future studies beyond the scope of the present proposal will investigate the correlation between viral binding and replication, combining *in vivo* experimental infections with *ex vivo* biologically relevant models to better disentangle viral-host interactions at the molecular level.

In conclusion, our results mapped the expression of α-2,3 and α-2,6 receptors in all species examined (sheep, goats, and alpacas). A prototype recombinant H5N1 virus of the 2.3.4.4b clade efficiently bound to the conjunctiva and respiratory tissues of the three species. Sheep tissues exhibited the highest binding levels across all tissues compared with goat and alpaca, whereas alpaca showed little to no binding to most tissues tested. More importantly, binding to tissues of minor livestock species is not a recent acquisition, as a prototype progenitor of the Gs/Gd lineage showed extensive binding in respiratory tract tissues. Similar results were obtained with pdmH1N1 and swine influenza virus strains. These findings suggest that factors beyond receptor-binding specificity play a pivotal role in the host range of H5N1 and that the role of minor livestock species in IAV ecology should be examined closely. Further studies examining the viral-host interactions of IAV, including H5N1 and others, in vivo and ex vivo will pave the way for a better understanding of IAV pathogenesis and the development of novel mitigation strategies.

## MATERIALS AND METHODS

### Institutional approvals and tissue acquisition

The Institutional Animal Care and Use Committee of Iowa State University approved this study (IACUC-24-090). The institutional biosafety committee approved laboratory work. Animals (n=3) of each species were procured from the Iowa State University Research Farms (sheep, goats) or from C&M Acres fiber mill (Maxwell, IA, USA; alpacas). Following euthanasia using Euthasol (Virbac AH, Inc., Fort Worth, TX, USA) intravenously, sections of the gastrointestinal and respiratory tracts were collected from these animals and were formalin-fixed and paraffin-embedded at the Veterinary Pathology Department at Iowa State University. The tissues collected from these animals included the conjunctiva, nasal turbinate, trachea, lung bifurcation, internal lung (including bronchi and alveoli), esophagus, and rumen.

### Cell culture

Madin-Darby canine kidney (MDCK) cells were kindly provided by Dr. Daniel R. Perez (University of Georgia, GA, US). Human embryonic kidney 293T (HEK293T) cells were purchased from ATCC (ATCC® CRL-3216^™^, Manassas, VA, USA). Cells were maintained in Dulbecco’s Modified Eagle’s Medium (Sigma-Aldrich, St. Louis, MO, USA) supplemented with 10% fetal bovine serum (FBS; Sigma-Aldrich, St. Louis, MO, USA), 1% Corning penicillin-streptomycin solution (AB/AM) (VWR, Manassas, VA, USA), and 1% L-Glutamine (Sigma-Aldrich, St. Louis, MO, USA). Cells were cultured at 37°C under 5% CO2. Cells were regularly screened for Mycoplasma using a Universal Mycoplasma Detection Kit (ATCC, Manassas, VA, USA) and were confirmed negative throughout the experiments.

### Reverse genetic plasmids

Reverse genetic (RG) plasmids encoding the 6 internal genes (PB2, PB1, PA, NP, M, and NS) of a laboratory-adapted version of A/Puerto/Rico/8/34 (H1N1)(PR8) and the full set of RG plasmids to obtain A/California/04/09 (H1N1) (pdm09 (H1N1)) were kindly provided by Dr. Daniel R. Perez (12, 65, 66). The low pathogenic (LP) version of the HA from A/turkey/Minnesota/22-010654-002/2022 (H5N1) clade 2.3.4.4b (LP-tk/MN/22 (H5N1)) and A/goose/Guan<u>g</u>dong/1/1996 (H5N1; ancestral) (LP-gs/GD/96 (H5N1)) were generated as previously described (24). Synthetic DNA encoding the LP-tk/MN/22 (H5N1) and the LP-gs/GD/96 (H5N1) HA and NA genes were procured from Twist Biosciences (San Francisco, California) and cloned into pHW2000 (1, 24). The generation of the DNA fragments for the RG plasmids for A/swine/Minnesota/A02245397/2020 (H1N1) (swH1N1) and A/swine/Iowa/A022861497/2023 (H3N2)(swH3N2) was carried out similarly and cloned into pJC2025 (ambisense plasmid like pHW2000) or pHW2000, respectively. All plasmid sequences were confirmed by Sanger sequencing (ISU DNA facility, Ames, IA, USA) and whole-plasmid sequencing (Psomagen, Inc., Rockville, MD, USA).

### Viruses

LP-tk/MN/22 (H5N1) and LP-gs/GD/96 (H5N1) were obtained by combining the respective HAs and NAs with the PR8 internal genes as previously described (12, 24). SwH1N1, SwH3N2, and pdm09 (H1N1) were obtained with the 8 segments of each virus. In all cases, reverse genetics experiments were performed as previously described (66, 67). Briefly, co-cultures of 9 ×10^5^ HEK293T and 1.5 ×10^5^ MDCK cells were seeded per well in a 6-well plate using Gibco Opti-MEM (Thermo Fisher Scientific, Grand Island, NY, USA) supplemented with 1% AB/AM. The following day, 1 µg of each plasmid was mixed with 18 µL of TransIT-LT1 transfection reagent (Mirus Bio LLC, Madison, WI, USA). The mixture was incubated for 45 min and then used to overlay the 293T/MDCK cells overnight. The next day, the transfection mixture was replaced with fresh Opti-MEM media containing 1% AB/AM. Twenty-four hours post-transfection, the media were supplemented with 1 µg/mL of tosylsulfonyl phenylalanyl chloromethyl ketone (TPCK) treated trypsin (Worthington Biochemicals, Lakewood, NJ, USA), and the same process was repeated at 72 hours post-transfection. Transfection supernatants were harvested at 96 hours post-transfection. Viral stocks of LP-tk/MN/22 (H5N1) and LP-gs/GD/96 (H5N1) were generated in 10-day-old specific pathogen-free (SPF) eggs (NVSL, Ames, IA, USA), while stocks of sw/MN/20 (H1N1) and sw/IA/23 (H3N2) were generated in MDCK cells. Allantoic fluids or cell culture supernatants were harvested at 48 or 72 h post-infection (hpi), respectively, centrifuged, aliquoted, and stored at −80 °C. Viruses were titrated by tissue culture infectious dose 50 (TCID50), and virus titers were established by the Reed and Muench method (68).

### *In vitro* growth kinetics

MDCK cells were seeded in a 6-well plate at a density of 400,000 cells/well in Gibco Opti-MEM supplemented with 1% AB/AM and incubated at 37 °C under 5% CO2 overnight. The next day, cells were inoculated with LP-tk/MN/22 (H5N1) and LP-gs/GD/96 (H5N1) at an MOI of 0.01, and the inoculum was incubated for 1 h at 37 °C. Next, the inoculum was removed, cells were washed twice with 1X PBS (Fisher Scientific, Fair Lawn, NJ, USA), and 3 mL of Opti-MEM containing 1 μg/mL TPCK-treated trypsin (Worthington Biochemical Corporation) was added to each well. Cell supernatant aliquots of 200 µl were collected at 0, 12, 24, 48, and 72 h post-infection and stored at –80 °C until further processing. Fresh OPTI-MEM supplemented with TPCK and 1% AB/AM was used to replenish the volume of media removed each time. To quantify viral replication, viral RNA was extracted using the MagMax-96 viral RNA isolation kit (Thermo Fisher Scientific, Carlsbad, CA, USA) according to the manufacturer’s instructions. Purified RNA was used as a template for RT-qPCR using the Quantabio qScript XLT One-Step RT-qPCR ToughMix kit (Quantabio, Beverly, MA) and specific primers targeting the M segment of the virus the FAM reporter and the TAMRA quencher were selected, and the cycles were as follows RT at 50°C for 20 minutes, RT deactivation at 95°C for 1 minutes and then for 35 cycles of denaturation at 95°C for 10 seconds and annealing at 60°C for 1 minute, after this results were opened and analyzed in the Quant Studio Design and Analysis software (Thermo Fisher Scientific, USA). Influenza viral titers were determined by comparison to a TCID_50_eq standard curve for an exact virus match with a known titer previously determined using the Reed and Muench method.

### Solid-phase binding assay

Each well in a ninety-six-well plate was covered by dispensing 0.2 mL of a fetuin working solution containing 1X coating buffer (Surmodics, Eden Prairie, MN, USA) and 10 μg/mL of fetuin (Sigma-Aldrich, St. Louis, MO, USA). Plates were then covered and incubated at 4°C overnight in a humidified chamber. The next day, plates were washed three times with DI water, then dried for 3 hours before being stored in a moisture-proof bag at 4°C. Each plate was incubated with 128 HA units/50 μL of the virus of interest in replicate columns and incubated at 4°C overnight in a humidified chamber. The next day, plates were washed 3 times with cold 0.02% Tween-80 (Thermo Fisher Scientific, Rockford, IL, USA) in PBS (washing buffer), and virus-coated plates were blocked with 0.1% desialylated BSA (BSA-NA) in PBS (blocking solution) for 2 hours at RT. The blocking solution (BSA-NA) contains 5% Bovine serum albumin (BSA) (Sigma-Aldrich, St. Louis, MO, USA) in PBS, PEN-STREP (Sigma-Aldrich, St. Louis, MO, USA), and 1 unit of neuraminidase sialidase from Vibrio cholerae (Sigma-Aldrich, St. Louis, MO, USA). The solution was incubated for 24 h at 37°C and then for 24 h at 60°C. After incubation with the blocking solution, the plates were washed twice, and 50 μL per well of 3’ SLN-PAA-biot (Biosynth, Bratislava, Slovakia) and 6’ SLN-PAA-biot (Biosynth, Bratislava, Slovakia) in serial 2-fold dilutions (Range:10 - 0.078 μg/mL) in reaction buffer (0.02% Tween-80, 0.1% BSA-NA, and 2 μM neuraminidase inhibitor (Toronto Research Chemicals)) were added and incubated at 4°C for 1 hr. Next, the plates were washed 5 times and incubated with 50 μL per well of HRP-labeled Streptavidin (Thermo Fisher Scientific, Rockford, IL, USA), diluted 1:1000 in Reaction Buffer, for 1 hour at 4°C. Plates were subsequently washed five times, and 50 μL per well of room-temperature TMB substrate (Thermo Fisher Scientific, Waltham, MA, USA) was added and incubated for 10 minutes at room temperature in the dark. Finally, the reaction was stopped with 50 μL per well of stop solution (Thermo Fisher Scientific, Rockford, IL, USA). The peroxidase activity (readout of viral binding) was determined at 450 nm using an Agilent BioTek Cytation 5 microplate reader. Wells without virus were used as negative controls, and the values in those wells were subtracted from the absorbance values from the virus-coated plates (69, 70).

### Viruses’ whole-genome sequencing

Nanopore-based next generation sequencing was carried out as previously reported (71). Briefly, viruses were sequenced on a MinION Mk1D nanopore sequencer using a native barcoding kit (SQK-NBD114-96). Sequencing libraries for each run were prepared following Oxford Nanopore Technologies (ONT) Influenza virus sequencing from RNA using SQK-NBD114 protocol (INF_9189_v114_revI_20Nov2024), following the manufacturer’s instructions. After library preparation the sequencing libraries were pooled and loaded onto a MinION R10.4.1 (FLO-MIN114) flow cell (ONT). The flowcell was checked before sequencing and run parameters were set for 16 hours. Basecalling was performed using Dorado High Accuracy v5.2.0 in MinKNOW 25.09.16. Reads with a Q score below 9 were filtered out. The generated FASTQ files were analyzed using EPI2ME v5.2.3 with the Flu pipeline (wf-flu) v1.3.1 from epi2me-labs (Oxford Nanopore Technologies), and the final consensus genome segments were generated from the resulting analyses.

### Lectin Histochemistry

The conjunctiva, nasal turbinate, trachea, lung bifurcation, internal bronchi of the lungs, tonsil, esophagus, and rumen from sheep, goats, and alpacas were used to determine the SA distribution using a lectin-based IHC assay as previously described (24). Briefly, formalin-fixed, paraffin-embedded tissue sections (4 µm) were mounted on Superfrost Plus slides (VWR International). After drying the slides at 60°C for 20 min, deparaffinization and staining were performed on the Roche Diagnostics Ventana DISCOVERY ULTRA research platform. The heat-induced epitope was retrieved using Cell Conditioning CC1 solution at 100°C for 24 min (Roche Diagnostics Corporation). Endogenous peroxidase was quenched for chromogenic staining using Discovery Inhibitor (Roche Diagnostics Corporation). Slides were then blocked with 1× Carbo-Free blocking solution (Vector Laboratories Inc.) for 32 min, followed by a Streptavidin/Biotin Blocking Kit (Vector Laboratories Inc.), with separate applications for 12 min each. Sections were incubated for 1 h at room temperature (RT) with one of the 3 lectins employed (*Sambucus nigra* lectin [SNA], *Maackia amurensis* lectin-I [MAL-1], *Maackia amurensis* lectin-II [MAL-2]; Vector Laboratories Inc.) at the listed concentration in Table 1. For SA α2,6, slides were incubated with biotinylated SNA, while biotinylated MAL-I/MAL-II were used for SA α2,3. SAs α2,6 and α2,3 was detected separately. Chromogenic staining was achieved using ready-to-use streptavidin-horseradish peroxidase (Vector Laboratories Inc.), which was incubated for 40 min on all 3 biotinylated lectins. The chromogenic signal was amplified with DISCOVERY Amplification anti-HQ Multimer (Roche Diagnostics Corporation). Chromogenic counterstaining with hematoxylin and bluing reagents (Roche Diagnostics Corporation) was performed, and cover slipping with Sakura Tissue-Tek film was performed. Negative assay controls omitted the primary lectin. Positive lectin staining was validated using bovine mammary tissues, as previously established.

Images and slides were observed and analyzed using an Olympus BX-43 light microscope equipped with an Olympus DP28 camera and CellSyns Dimension version 4.1.1. At least 4 board-certified veterinary pathologists from the American College of Veterinary Pathologists evaluated all tissues, and results were reported as present or absent using qualitative terms. Each receptor was assessed on a scale from 0 to 4, with 0 indicating no positive labeling throughout the examined section, 1 indicating weak positive labeling (approximately 1-5% of the examined sections), 2 indicating mild positive labeling (approximately 6-20% of the examined sections), 3 indicating moderate positive labeling (approximately 21-50% of the examined section), and 4 indicating strong positive labeling (>51% of the examined section). Localization (Table 2) indicates where positive labeling was most prominent, either on the surface of epithelial cells of the examined tissue (SE) or within goblet cells (GC). In instances where no tissue was present in the examined sections, the tissue was not scored, and the result was indicated with a hyphen (−)

### Virus-binding assay

For deparaffinization, the slides were placed in the Epredia™ Coplin Staining Jar and heated at 60°C for 1 hour. Then the slides were placed in Citrisolv Xylene (VWR, USA) for 15 minutes, followed by 100% Ethanol (EtOH) for 2 minutes twice in a different container each time, and finally in 95% EtOH for 2 minutes twice in a different container each time. The slides were washed in tap water, then in DI water for 5 minutes. For antigen retrieval (Heat-induced epitope retrieval, HIER), the slides were placed in a staining jar, immersed in 1X citrate buffer (Abcam, Cambridge, UK) at pH 6, and incubated at 95°C for 45 min. The slides were then cooled for 1 hour at room temperature and washed twice with PBS for 2 min each, and once for 10 min. The slides were incubated overnight with the virus of interest, standardized to 100 HA units/50 μL diluted in PBS. The HA units of each virus were standardized using a solution of 0.5% turkey red blood cells. Slides were incubated with clean allantoic fluid as a negative control. After virus incubation, all slides were treated with 10% formalin solution for 30 seconds and washed twice with PBS. Slides were then incubated with Bloxall (Vector Laboratories, Newark, CA, USA) to block nonspecific staining caused by endogenous HRPs for 10 min at room temperature. Then wash twice with PBS for 5 min. Afterward, 5% BSA (Sigma-Aldrich, St. Louis, MO, USA) was used to block for 1 hour and the slides were washed twice with PBS. For primary antibody incubation, an anti-influenza A virus HA (Clone CR9114, Cell Sciences, Newburyport, MA, USA) was used at a 1:500 dilution in PBS-BSA 3% overnight at 4°C. The next day, the slides were washed 3 times with PBS and once with Tris-buffered saline-Tween 20. For the secondary antibody incubation, a Rabbit anti-Human IgG secondary antibody-HRP (Thermo Fisher Scientific, Waltham, MA) was used at a 1:1000 concentration for 1 hour at room temperature. After incubation, slides were washed 3 times with PBS for 5 min each. Finally, diaminobenzidine substrate (Cell Signaling, Danvers, MA,USA) was applied to the tissues for 3 minutes, then the reaction was stopped with DI water wash for 3 minutes. Finally, the slides were immersed in Mayer’s Hematoxylin (Sigma-Aldrich, St. Louis, MO, USA) for 30 sec., and the reaction was stopped with tap water. To dehydrate the slides, they were placed in staining jars containing: 1) 95% EtOH 3 times for 1 minute, 2) 100% EtOH 3 times for 1 minute, and 3) Citrisolv twice for 3minutes. The slides were mounted with mounting medium and a coverslip was applied. Images and slides were observed and analyzed using an Olympus BX-43 light microscope equipped with an Olympus DP28 camera and CellSyns Dimension version 4.1.1. The virus-binding in each tissue was assessed on a scale from 0 to 4, with 0 indicating no positive labeling throughout the examined section, 1 indicating weak positive labeling (approximately 1-5% of the examined sections), 2 indicating mild positive labeling (approximately 6-20% of the examined sections), 3 indicating moderate positive labeling (approximately 21-50% of the examined section), and 4 indicating strong positive labeling (>51% of the examined section). Localization (Table 2) indicates where positive labeling was most prominent, either on the surface of epithelial cells of the examined tissue (SE) or within goblet cells (GC). In instances where no tissue was present in the examined sections, the tissue was not scored, and the result was indicated with a hyphen (−)

### Graphs and panels

All data analyses and graphs were performed using GraphPad Prism software version 11.0.2 (100) (GraphPad Software Inc., San Diego, CA, USA). Panel figures and schematic representation of the viruses were generated in BioRender as previously described (72). Illustrations from Figure 6 were obtained from the NIH Bioart repository.

## AUTHOR CONTRIBUTIONS

NM, EB, CJC, SC, and TMB designed the experiments. TMB, BB, and OF obtained the tissues used in this study. NM, SO, GB, and CJC performed the reverse-genetic assays and characterized the viruses. JGT and RP performed the lectin assays and generation of the H&E slides for further analysis. LB, AH, and MZ performed viral next-generation sequencing. NM, EB, and GB performed the virus-binding assays. TMB, EB, SC, and EC performed scoring of the different tissues. NM, EB, RN, OF, EC, SC, TMB, and CJC performed data analysis and discussion. NM and CJC wrote the first manuscript draft. NM, EB, SC, TMB, and CJC edited the manuscript. SC, TMB, and CJC provided funding. All authors approved the manuscript.

## ACKNOWLEDGEMENTS

We are grateful to the Veterinary Pathology Department for assistance with necropsies and to the technical staff for facilitating the acquisition of animals, the performance of necropsies, and the preparation of histologic slides. We also thank Amanda La Coco, Olivia Stewart, Matt Brewer, and Katy Martin for their technical support, as well as Dean Isaacson and the personnel of Laboratory Animal Resources for their assistance with animal acquisition and care. We further acknowledge the personnel of the Veterinary Pathology Department and Veterinary Diagnostic Laboratory for their assistance with tissue processing and preparation of the necessary slides. This work was partially funded through the Iowa Livestock Health Advisory Council (ILHAC) and start-up funds from Iowa State University to CJC. Financial and programmatic support was also provided by the Iowa State University College of Veterinary Medicine through the Mapes Wildlife Care, Education, and Research Endowment, as well as by start-up funds provided by Iowa State University to TMB. Additional funding includes resources from the vice president for research office to CJC and SC, and a research agreement from USDA to TMB, CJC, and SC (AP24VSD&B000C077).

